# Multi-decadal warming alters predator’s effect on prey community composition

**DOI:** 10.1101/2024.03.19.585501

**Authors:** Jingyao Niu, Magnus Huss, Aurélie Garnier, Anti Vasemägi, Anna Gårdmark

**Affiliations:** Swedish University of Agricultural Sciences, Department of Aquatic Resources, Box 7018, 750 07 Uppsala, Sweden; Swedish University of Agricultural Sciences, Department of Aquatic Resources, Institute of Freshwater Research, Stångholmsvägen 2, SE-178 93 Drottningholm, Sweden; Université de Rennes, UMR 6553 CNRS ECOBIO, 263 avenue du Général Leclerc, 35042 Rennes, France; Estonian University of Life Sciences, Institute of Veterinary Medicine and Animal Sciences, Chair of Aquaculture, 46A Kreutzwaldi St., 51006 Tartu, Estonia

**Keywords:** local adaptation, thermal evolution, climate warming, trophic interactions, *Perca fluviatilis*

## Abstract

Predator responses to warming can occur via phenotypic plasticity and evolutionary adaptation, resulting in changes in their prey communities. However, we lack evidence of how warming-induced evolutionary changes in predators can influence the food web. Here, we ask whether fish subject to long-term warming across multiple generations differ in their impacts on prey communities compared to their nearby conspecifics experiencing a natural thermal regime. We carried out a common garden mesocosm experiment with larval perch (*Perca fluviatilis*), originating from one heated or one reference coastal environment, feeding on zooplankton communities under a gradient of experimental temperatures. We found that fish thermal origin influenced the zooplankton communities, and differently so depending on the experimental temperature. In presence of fish of heated origin, there were less zooplankton and also fewer individuals of large size, except for at intermediate experimental temperatures. Our findings show that differences between fish populations, potentially representing adaptation to local thermal environment, caused by multi-generational warming can cascade down via trophic interactions to also affect their zooplankton prey communities. Considering climate warming, our results suggest that rapid evolution in predators might have indirect cross-generational ecological consequences propagating through food webs.

## Introduction

Organisms respond to prevailing climate warming by means of plasticity during acclimatization and evolutionary changes over generations (1). Despite extensive efforts in investigating species-(2) and community-responses to warming (3,4), such studies are mostly on ecological time scales (5–8). When studies do span multiple generations, they are rarely field-based but instead under laboratory conditions (9,10) or the thermal gradient involved in the study encompasses a large geographic area including confounding factors (11). Yet, given fast ongoing environmental changes due to global warming, there is pressing need to investigate the cascading effects of potential adaptations to warming in natural predators on prey communities, to improve our ability to predict the impacts of global change on food web stability and ecosystem functioning (12).

Ectotherm individual physiological responses to warming, such as an increase in metabolic rate (13–15), can affect organisms’ activity level, feeding and locomotion (16–18). Physiological responses to warming can ultimately result in faster growth at younger life stages (19,20), reduced size at maturation (21) and lowered survival (22). Warming-induced changes in predator feeding rate and behaviour can, in turn, lead to reduced biomass, shifts in species composition and size distribution (23), as well as behavioural and morphological changes in their prey communities (24), which can have profound impact on the ecosystem (25). In aquatic systems, fish commonly exert top-down control on zooplankton communities by direct size-specific removal of individuals with potential effects on prey population dynamics and viability (26). However, how long-term warming-induced responses in fish predators affect prey communities is largely unknown.

How do fish adapt or acclimatize to warming in ways that may affect their prey? Fish responses to multi-generational warming include changes in their metabolism, growth, development, morphology and body size (14,20,27,28) which could directly or indirectly affect predators’ feeding. First, increased metabolic rate is common in warmer environments (29), which requires and also enables organisms to consume more prey (30). In case they cannot fully compensate by increased feeding rate, increased metabolic cost can result in lower net energetic gains in warmer environments, with less energy being allocated to activities such as growth, locomotion and even motivation to initiate predation (31,32).

However, to counter the increased energetic deficiency in a warmer environment – a compensatory metabolic response – a depression in standard metabolic rate (SMR) has been found in several warm-acclimatized or warm-adapted fish species (14,33–35). A lower SMR would reduce starvation risk, as individuals require less food to maintain their energy balances. Compensatory metabolic responses can also lead to changes in locomotor performance that directly enhance feeding behaviour (5). Another indirect way by which adaptation to warming may lead to increased food intake is through faster development (36) or body growth (10) in warm environments. A faster early growth can lead to a larger body size when young, thus potentially a larger gape size (37) or a better body condition at a given age that can lead to an advanced switch in diet towards larger prey (10,38,39). Changes in development or growth can also cause phenological shifts among predators and prey. For example, shifting relative size distribution or turnover rates can result in predator-prey mismatches (38). In cases where prey communities have adapted to warming, predator fish developed feeding morphological changes in response to changes in their food availability and composition (27). Following this logic, prey should be affected by warming-induced long-term changes of their predators, as suggested by changes in trophic interactions in short-term warming acclimatization experiments (5,7) and aquatic invertebrates (11,40). However, responses in prey communities to such indirect effects of multi-generational warming via trophic interactions are yet to be demonstrated.

Effects on prey communities from warming-induced responses in predators can play out differently depending on the extent of temperature change relative to their natal habitat temperature, and exposure time. Effects of adaptation or acclimatization on, for example, thermal tolerance or SMR often follow a hump-shape function of the novel environmental temperature (14,41). Therefore, the indirect effect from these responses on predator feeding can also be hump-shaped, which has been shown for attack and maximum ingestion rates (5,18,42,43). While there is evidence for thermal adaptation in predators (14) and for that thermal acclimatization in predators can shape predator-prey interactions (5,7), we still lack experimental tests of how cross-generational responses in predators to warming may influence prey community composition under different temperatures.

Here, we investigate whether the thermal origin of larval fish affects zooplankton prey communities, and if this effect varies with experimental temperature. To answer these questions, we carried out a common garden mesocosm experiment using Eurasian perch (*Perca fluviatilis*) larvae hatched from eggs collected in two adjacent wild populations inhabiting ecosystems with different thermal regimes: heated or unheated for 41 years. The two neighbouring coastal areas share similar environmental characteristics, except for the 41 years of +5-10 °C difference in water temperatures that spans across multiple fish generations (Figure S1). We tested the effect of fish larvae of different thermal origin on a natural (from an unheated area) zooplankton prey community and manipulated the mesocosm water temperatures, rendering a gradient of mean temperatures from 14 to 25 °C over the experiment. This allowed us to study the potential for effects of warming across trophic levels based on a wild, potentially warm-adapted, fish population exposed to multi-generational heating, feeding on a natural prey community. Our hypotheses are: 1) changes in experimental zooplankton communities depend on the thermal origin of the fish larvae feeding on them, and that 2) these differences vary with temperature. Our findings imply that impacts of climate warming on predators across generations can have indirect, yet substantial, effects on their prey communities via shifts in trophic interactions.

## Methods and materials

### Fish larvae thermal-origin

The larval Eurasian perch (*Perca fluviatilis*, hereafter perch) used in the experiment originated from parental populations from areas that either has undergone heating for 41 years or is a control population residing in an adjacent coastal area experiencing a natural temperature regime. The heated population has been residing in an 1-km^2^ heated area since 1980 and has been subjected to a temperature of 5-10 °C above the natural, caused by warm water discharge from nearby nuclear reactors on the western Baltic Sea coast (60.43N 18.19E, Figure S1). The design of the set-up ensured that the two populations have been exposed to similar environmental conditions, other than the heated water flow-through (i.e. the two adjacent areas share the same coastal environmental characteristics and water properties, see Supplement: Perch larvae thermal origin). In the heated area, perch have larger female size at age and higher growth rates when small (20) but slower growth in large males (28), higher mortality (22), more advanced gonad development at a given time of year (44), and a smaller female maturation size (21). When experimentally exposed to acute warming, large perch from the heated area displayed lower metabolic rate, thermal sensitivity and higher thermal plasticity than perch from the reference area (35). Allelic composition shifts in the MHC alleles have been observed in the heated-area perch population (45), which might explain the higher parasite resistance in the heated population (46). Perch in the heated area have thus separated from the perch population in the adjacent unheated control area both genetically (45,47) and phenotypically (20,21,28). However, the metal grid that prevented the exchange of organisms between the areas was removed in 2004. The grid removal increased the probability of larger fish (>10 cm, 48) dispersing between the areas, although the strong current (∼100 m^3^/s) likely prevents immigration of small or poorly swimming individuals into the heated area.

To study the effects on prey communities of larval perch from these two geographically close but thermally contrasting habitats, we collected 15 separate roe strands (Table S1) from each area on 2021 May 10^th^. Eight strands with similar width and adhesion between eggs were selected out of 15 from each habitat and transferred to indoor aquaria for hatching. Each egg strand was placed in separate aquaria with ∼ 100 L aerated, unfiltered coastal water with a 16 h light: 8 h dark photoperiod indoors at room temperature. The rest of the roe strands were kept in smaller aquaria (∼40 L) for genetic analysis. Perch larvae started hatching from May 17^th^, and on May 22^nd^ there was an adequate number of larvae to use for the experiment (for hatching record, see Table S1). We selected two newly hatched (< 5 days old) perch larvae from each of five egg strands per origin (heated or unheated area) based on similar hatching time and egg strand widths (Table S1). At the start of the experiment, we acclimatized the fish to the experimental mesocosm by gradually lowering and immersing the bag containing aquaria water and larvae into the mesocosm water. Besides introducing perch larvae to the mesocosms, we also sampled larvae (or egg, if no larvae were hatched) from all 15 roe strands per origin on the same day for larval body size measurements and/or genetic analyses. Our microsatellite DNA analysis based on 14 loci (as described in supplementary material 14_microsatellite_primer.xlsx), using three individuals each from these 30 roe strands (supplementary material microsatelliteDNA_genotypes.xlsx), showed low but statistically significant genetic differentiation between the heated and unheated individuals (Fisher’s exact probability test, Chi^2^ = 82.5, P < 0.001; Fst = 0.006).

### Mesocosm experiment

We established 38 outdoor mesocosms using tanks (FlexiTank, Max Grow Shop) of volume ∼400 L, diameter 0.68 m, height ∼1.1 m, made of polyurethane fabric and supported by rods. Altogether, 26 of these tanks were inoculated with fish larvae (10 individuals from one of the two origins per tank) and the remaining 12 tanks were kept without fish as control (Figure S2, Table S2). The experiment was conducted outdoors for 20 days (2021 May 22^nd^ – June 10^th^, experiment day 0-20). In these open-to-atmosphere tanks, we generated a simplified pelagic food web with plankton and fish larvae. The mesocosms were filled with 350 L water from the unheated area (salinity ∼5 PSU) filtered through 50 μm mesh. Water filling underwent from May 12^th^ – 18^th^, ensuring that approximately equal amounts of water were added to all mesocosms each day. One day prior to the start of the experiment (i.e. on May 21^st^), zooplankton communities were collected at four sites in bays and along the shore in the unheated area, partly overlapping with the site for egg strand collection. One plankton net of 20 and one of 70 μm mesh size were lowered to the target depth (< 1 m), pulled horizontally for approximately 5 min at average speed of 1 knot (0.5 m/s), and thereafter quickly retrieved to the surface to empty collected plankton. Active sampling was conducted for in total three hours. Collected plankton were kept in one ∼700 L tank filled with filtered water from the unheated area until being added to the mesocosms. 7.5 L of this well-mixed plankton mixture was added to each mesocosm on May 21^st^.

Experimental temperature in each mesocosm was manipulated by thermostats (Eheim 300W) placed in the centre of the mesocosm for heating, or by cooling with a surrounding flow-through system (Figure S2). A thermal gradient of 14 - 25 °C was maintained among the 38 mesocosms throughout the course of the experiment, with natural variation due to weather and time of day (Figure S3). We deployed a temperature logger (HOBO UA-001-64 Pendant Temperature Data Logger) right after temperature manipulation started at 40 cm depth in each mesocosm to record temperature every hour. Each mesocosm was aerated on the bottom with approximately the same air flow intensity using one air stone connected to an air pump (Airflow 400, IP44 230V). The aeration saturated mesocosm water with oxygen and prevented temperature stratification. Note that while dissolved oxygen concentration declined slightly with temperature (a physical law that applies to all systems), the percentage saturation was independent of temperature across the gradient (> 100%).

### Sampling and sample processing

Zooplankton and chlorophyll *a* (chl *a*) were sampled prior to the addition of fish larvae to the mesocosms (on day 0 and day −1, respectively), in the middle of the experiment (day 9) and when approaching the end of the experiment (day 19) by sampling 3.3 L water from each mesocosm at 40 cm depth using a 0.66 L Ruttner water sampler. The water was filtered through a 70 μm mesh to keep most zooplankton on the mesh and reduce the amount of water. We gently rinsed off each sample of the zooplankton collected on the mesh into a 100 ml brown bottle with tap water and added 4 ml Lugol’s iodine solution into the same bottle. To estimate chl *a*, we filtered 500 ml water through a 47 mm glass fiber filter (Whatman^TM^ GF/F) using a pressurized chamber. The glass fiber filters were stored folded in aluminum foil and put in sealed bags at −20 °C until they were processed for estimation of chl *a* concentration to approximate the phytoplankton biomass in our mesocosms (49). Before adding perch larvae to the mesocosms, chl *a* and zooplankton were sampled the first time to confirm that there were no significant differences in phyto- and zooplankton biomass and zooplankton composition among mesocosms of different treatments (Supplement: Mesocosm initial conditions). From June 3^rd^, we put the remaining filtered water after sampling back to mesocosms to slow down the noticeable decrease in water level (mostly due to evaporation under warm weather).

Zooplankton identification, counting and measurements were done using a stereo microscope (Leica M125C). We counted and measured individuals in 30 ml subsamples of the zooplankton mixture, and in the remaining volume continued counting only the taxa of which less than 50 individuals were counted in the first 30 ml. *Copepoda* were separated into nauplii, copepodite stage 1-3, stages 4-5 of order *Calanoida* (genus *Acartia* or *Eurytemora*), adults and order *Cyclopoida*. Cladocerans were separated into *Bosmina* sp, *Chydorus* sp and *Podon* sp. Rotifers were identified to genus *Keratella* (*K. cochlearis, K. quadrata and K. cruciformis*), *Bdelloida* and *Brachionus*. We measured the individual body length to the nearest 0.01 mm of 20 or more individuals in each taxon for each mesocosm. Zooplankton biomass was estimated from length measurements using body length-biomass relationships (Table S3).

To estimate chl *a* concentrations, we processed the glass fibre filters as follows: we cut each filter in half (exact proportion measured for accuracy) and put one of the halves directly into a 5 ml screw cap vial filled with 96% ethanol and then kept in darkness at 4°C for 22.5 h for chl *a* extraction. All samples were shaken vigorously halfway through the extraction. Samples were thereafter centrifuged at 5000 rpm for 5 min to sediment any particles. Three replicates of 200µl of the supernatant of each sample were pipetted into three wells of a black solid-bottom 96-well plate. Fluorescence was measured at λex/em = 444±12/675±50 nm using a microplate reader (Hidex Sense) and converted to chl *a* concentration (μg/L) following Equation S1. The samples and plates were kept cool in darkness and handled as fast as possible during the process.

Perch larvae were sampled at the end of the experiment, on day 20, by hand-made drop nets of 1 mm mesh size that were slightly smaller than the mesocosm tank’s diameter. We sank the drop net vertically to the mesocosm bottom, turned it horizontally at the bottom, pulled to the surface horizontally and picked up the fish from the net using tweezers. The fish were immediately put in a benzocaine solution (200 mg/L) to euthanize them and then transferred to containers with 80% ethanol for storage. If no fish were caught during five such fishing attempts, we deemed that no fish was left in the mesocosm and moved on to the next. In total, 138 fish larvae were caught. For the first three mesocosms, we repeated the same fishing process to validate the drop net method and no fish were caught during this process. However, 16 fish were found alive 22 days after the experiment ended (July 2^nd^) when we emptied the mesocosms. The number of fish captured on both occasions (day 20 and July 2^nd^) can be found in Table S2. Body lengths of caught perch larvae were measured to the nearest 0.01 mm using a stereo microscope (Supplementary material Table S4.xlsx). Both the standard length and total length were measured for the mesocosm-caught fish whereas only total length was measured for one-week-old larvae collected from the hatchery aquaria. Fish wet weights were measured to the closest 0.1 mg after we patted them dry on both sides twice to minimize alcohol residual (Table S4 & S5).

### Statistical analyses

We used analysis of variance (ANOVA) to test whether thermal origin of the larval fish, experimental temperature and their interaction had an effect on zooplankton abundance, biomass and composition (Table 1 & 2). We also tested for the effect of time (i.e. experiment day) in analyses where data encompassed the whole experiment, and the random effects of mesocosms (Table 2). Experimental temperature of each mesocosm was calculated by averaging the hourly measurements from day 0 up until the experiment day in question. We treated experimental temperature as a continuous variable as the temperatures measured were quite evenly distributed across the full gradient of 14 – 25 °C, and also fluctuated within each mesocosm due to exposure to natural temperature variation (Figure S3).

**Table 1.** Results of ANOVA on the effects of larvae thermal origin, temperature and the interaction between origin and temperature on zooplankton community abundance and biomass at the end (day 20) of the experiment. Symbol * indicates significant results (P < 0.05). Best model estimates are the coefficients from the best model based on model selection (Table 2).

| Zooplankton | Explanatory variables | F(1, 38) | P | Best model estimates |
| --- | --- | --- | --- | --- |
| **Total abundance** | Origin | 5.84 | 0.024 * | 5.58 |
|  | Temperature | 5.12 | 0.034 * | - 0.02 |
|  | Origin x temperature | 4,82 | 0.039 * | - 0.34 |
| **Total biomass** | Origin | 4.60 | 0.043 * | - 0.79 |
|  | Temperature | 2.33 | 0.141 | - 0.08 |
|  | Origin x temperature | 1.30 | 0.266 | / |

**Table 2.** Results of model selection regarding zooplankton abundance based on model pair testing - likelihood ratio tests (significance level between the new model and null model). Model pairs are: null and second, second and third, and so on. We judged whether including larvae origin, or experimental temperature and larval origin * temperature, improved the model explaining the total variance based on the significance of likelihood ratio tests. The best models’ (in bold) coefficients are origin (unheated) 5.58, temperature −0.02 and origin x temperature −0.34; origin (unheated): −0.47, temperature: −0.059.

|  | End of experiment |  | Throughout the experiment |  |
| --- | --- | --- | --- | --- |
|  | Formula | Significance | Formula | Significance |
| ln(total zooplankton abundance + 1) ~ | 1 |  | day + random(day\|mesocosm) |  |
|  | temperature | 0.07 | temperature + day + random(day\|mesocosm) | 0.07 |
|  | origin + temperature | 0.02* | **origin + temperature + day + random(day\|mesocosm)** | **0.02 \*** |
|  | **origin \* temperature** | **0.03\*** | origin * temperature + day + random(day\|mesocosm) | 0.14 |

We used a model selection approach to test for effects of each variable. Models were established using the R-packages lme4 (50) and betareg (for proportional data, 51). We set the null model as response ∼ temperature + day + random(mesocosm), where the random effect was only included when we analyse more than one day of data. For model selection, we ranked the models based on significance of pairwise likelihood ratio tests on sequential models of increasing complexity (52). Data visualization and processing were done using the packages within the tidyverse collection (53).

Zooplankton abundance, biomass and fish larvae weight were ln-transformed prior to analyses and fish survival rate (sum of larvae caught on day 20 and July 2^nd^, divided by 10) was square root transformed to account for heteroscedasticity. For taxon-specific analysis, we grouped Calanoid copepodite stage 4-5, adult *Eurytemora* and *Acartia* and Cyclopoid copepods together as *Copepoda* because the numbers were low (due to fish predation, see Supplement: Fish predation). Using the function vegdist in package ‘vegan’ (54), we calculated the distance matrix of Bray-Curtis dissimilarity indices (55) to represent zooplankton community composition based on abundance and on biomass. We then used the function capscale to conduct an Analysis of Principal Coordinates, capscale(matrix ∼ 1), on each of the distance matrices in two steps: (1) the dissimilarity matrix was ordinated using function cmdscale, and (2) the ordination was projected to maximize the variance in each ordination axis (56). We then used an ANOVA to test if the variance found among mesocosms, as represented by the first ordination axis (PCO1), differed depending on fish larval origins. We also recorded variance found between zooplankton taxa (species scores on PCO1), to investigate whether some taxa were particularly influential in driving the variation in zooplankton composition among mesocosms. For analysis of zooplankton size composition, we found that 200 μm was a suitable threshold to separately group large- and small-sized zooplankton for their clear difference in abundance. It also separated the zooplankton species known to be less preferred as prey by fish larvae (e.g. rotifers with lorica, a hard structure making them unfavorable prey for perch larvae, 57) from their preferred prey (more than 99% of *Keratella* were below 200 μm). We calculated the proportion of large-sized (> 200 μm) zooplankton and tested for effects of larval thermal origin, experimental temperature and the interaction using model selection as described above. Finally, we made predictions using the best model according to the model selection results.

All data processing and statistical analyses were conducted in R, version 4.0.2, R Core Team 2014.

## Results

Fish larvae origin had a significant effect on zooplankton total abundance as well as biomass at the end of the experiment (Table 1, Figure 1a & Figure S4), but differently so depending on the experimental temperature (Table 1 & 2). Indeed, no effect of temperature on zooplankton was found for treatments with larvae of heated origin, whereas temperature had a negative effect on zooplankton abundance in mesocosms with fish from the unheated area (Figure 1a; ANOVA on zooplankton difference at high temperatures, P = 0.024). The thermal origin of fish affected zooplankton total abundance throughout the experiment (Figure 1b, Table 2), but zooplankton total biomass only at the end of the experiment (day 19, Table S6). Moreover, the interaction between larvae origin and experimental temperature did not affect zooplankton abundance across the duration of the experiment but only became detectable at the end of the experiment, likely due to changes emerging between day 9 and 19 (Table 2). The effect of fish origin on zooplankton abundance and biomass was not due to differences in fish abundance nor their difference in growth as the number of larvae caught and their total length at the end of the experiment did not differ between origins (Figure S5a & S5c, ANOVA on survival: F(1,22) = 0.44, P = 0.48; length: F(1,134) = 2.58, P = 0.111).

**Figure 1.**
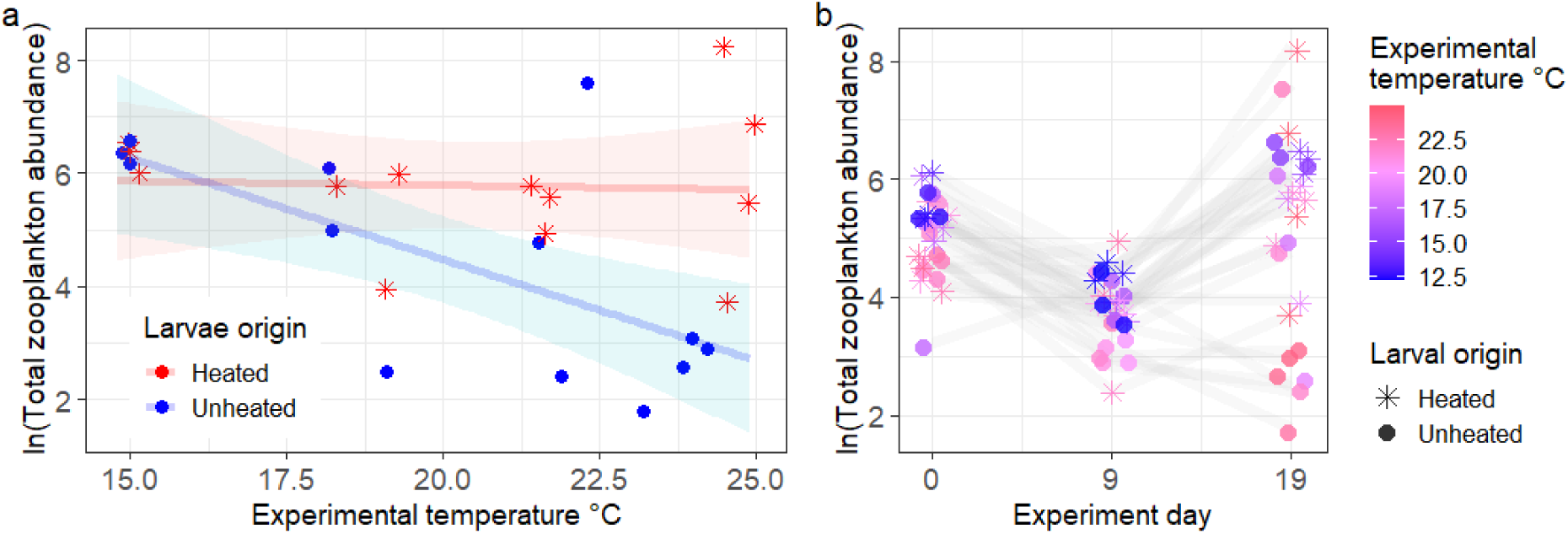
Zooplankton total abundance at the end of the experiment (a) and throughout the experiment (b) in mesocosms containing larval fish originating from the heated (stars) or unheated population (circles) at different experimental temperatures. The red and blue lines depict the predicted (from *ln(abundance) ∼ origin + temperature + day +random(day|mesocosm)*, Table 2) ln-transformed zooplankton abundance and the belts show their corresponding confidence intervals at 95%. The effect of larvae origin on zooplankton abundance is predicted to increase with temperature. Note that all mesocosms were measured on the same days (0, 9, 19), but have been displaced for visibility.

Zooplankton compositional variation based on abundance (Figure 2a) or biomass (Figure S6) as described by PCO1 significantly correlated with larval origin (Table 3, P < 0.05 for the term larval thermal origin, Figure 2b and Figure S6, η2 = 0.18). The effect of larval origin on zooplankton community composition varied with experimental temperature (Table 3, P < 0.05 for the interaction term origin * temperature, η2 = 0.10), with the greatest difference in zooplankton community composition between mesocosms containing larvae of heated versus unheated origin at high temperatures (Figure 2b & S7). The six taxa partitioned most variance on PCO1 (Figure 2a), which correlates significantly with fish larvae thermal origin (Table 3), were *Keratellas*, nauplii, copepodite and *Cyclopoid* copepods.

**Figure 2.**
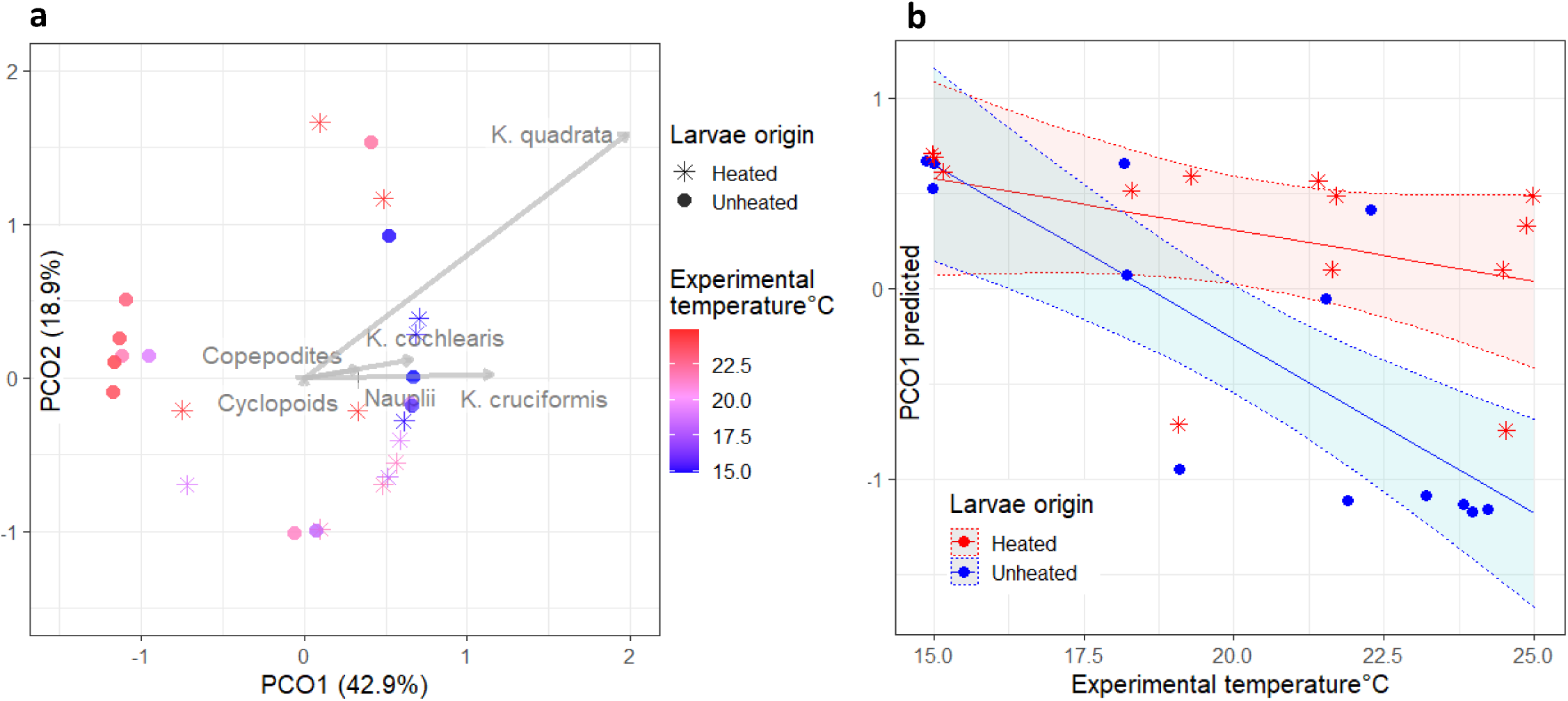
Ordination (a) showing PCO1 (explaining 42.9% of the total variation) and PCO2 (explaining 18.9%) results on zooplankton community composition based on abundance of different taxa in mesocosms with fish larvae from the heated (star) or unheated population (circles). Colour shows mean experimental temperature (from blue to red). Predicted PCO1 (b) at given experimental temperature, where colour indicates larvae origin (heated vs unheated). Predictions made from linear model *lm(PCO1 ∼ origin * temperature).* The difference in zooplankton composition between larvae origins increased with temperature.

**Table 3.**
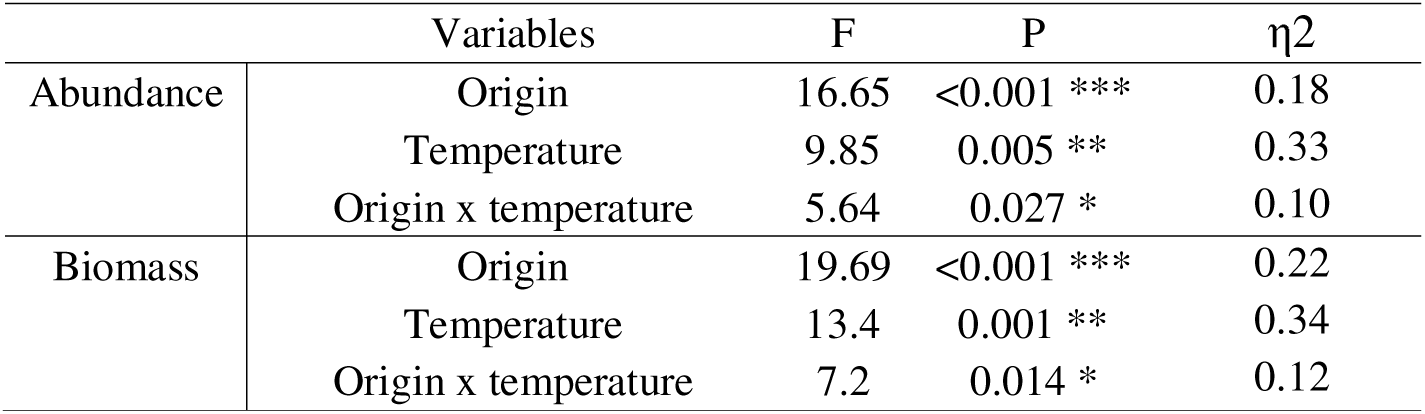
Results of an ANOVA model on the effects of fish larvae thermal origin (heated/unheated), experimental temperature and their interaction on PCO1 based on zooplankton community compositional abundance or biomass. We found no significant between-mesocosm differences in PCO2, based on abundance or biomass, in relation to fish origin or experimental temperature.

| | Variables | F | P | $\eta^2$ |
| --- | --- | --- | --- | --- |
| Abundance | Origin | 16.65 | <0.001 *** | 0.18 |
|  | Temperature | 9.85 | 0.005 ** | 0.33 |
|  | Origin x temperature | 5.64 | 0.027 * | 0.10 |
| Biomass | Origin | 19.69 | <0.001 *** | 0.22 |
|  | Temperature | 13.4 | 0.001 ** | 0.34 |
|  | Origin x temperature | 7.2 | 0.014 * | 0.12 |

The relative abundance of large-sized zooplankton differed depending on fish origin, but in different ways depending on the experimental temperature (Figure 3, interaction term origin * temperature P < 0.001 in Table S7). At higher temperatures (above 20 °C in Figure 3), there were significantly less large-sized zooplankton in mesocosms with larvae of heated than unheated origin (ANOVA, F(1, 12) = 6.07, P = 0.030), whereas the opposite was observed at low temperatures. Among zooplankton taxonomic groups, copepods constituted a significantly smaller proportion in mesocosms with larvae of heated than unheated origin (ANOVA, F(1, 12) = 5.78, P = 0.033). Larval thermal origin thus affected both zooplankton size- and species composition, as well as the relationship between zooplankton composition and experimental temperature.

**Figure 3.**
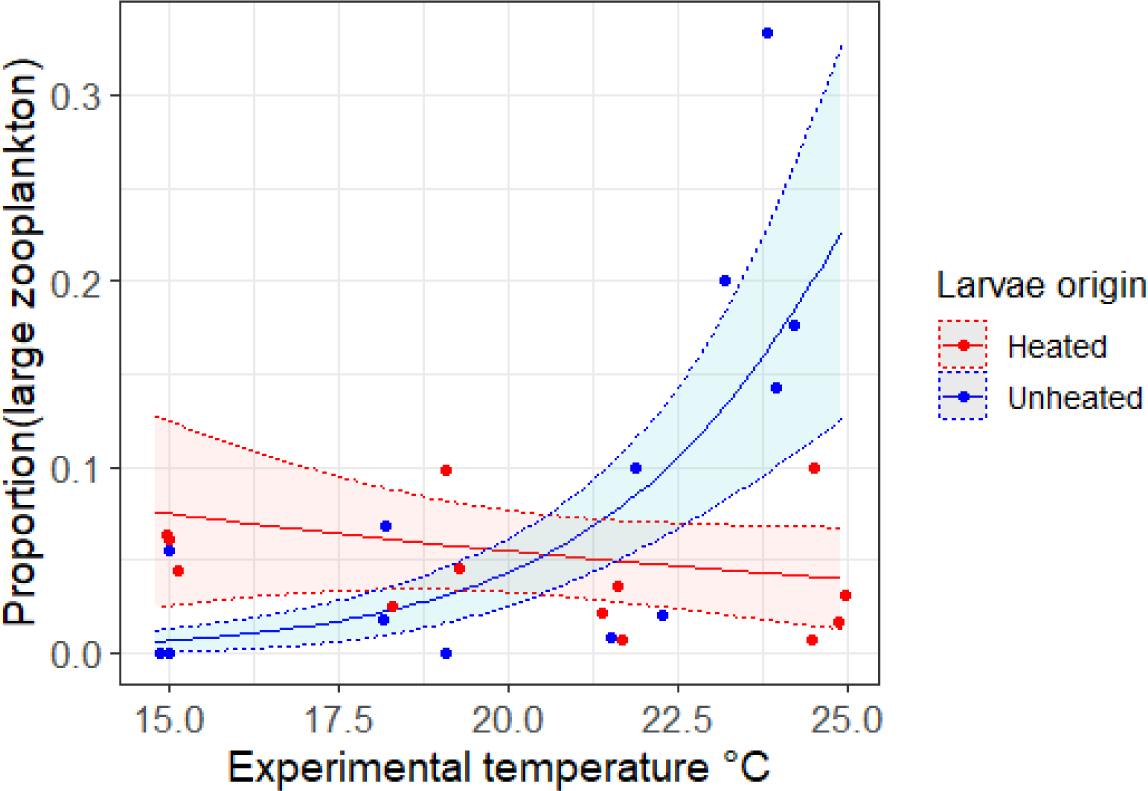
Proportion of large-sized zooplankton in mesocosms containing larval fish of heated (red) or unheated (blue) origin. Predictions made from the best model *P(large zooplankton) ∼ origin × temperature* (Table S7) show that fish origin had a clear effect on how fish from that origin affected large zooplankton under different experimental temperatures.

Unlike number of fish caught or individual body length of larvae, individual larvae of heated origin gained slightly more weight than the ones of unheated origin during the experiment (ANOVA, P = 0.038, η2= 0.03). Experimental temperature also increased individual weight of larvae independently of their thermal origin (Figure S5 b, ANOVA, F(1, 134) = 55.54, P < 0.0001, η2 = 0.28), which was also the case for body length (Figure S5 c, ANOVA, F(1, 134) = 41. 17, P < 0.0001, η2 = 0.23).

The differences in zooplankton abundance, biomass and composition between mesocosms with fish larvae of heated or unheated origin was likely not caused by any differences in phytoplankton biomass because chlorophyll *a* concentration did not differ between mesocosms containing perch larvae of heated or unheated origin (Figure S8a, ANOVA, P=0.462). However, without fish, chl *a* increased with temperature (Figure S8b, R^2^= 0.394, P < 0.0001).

## Discussion

We found that the thermal origin of fish larvae affected the community abundance, biomass and composition of their zooplankton prey in a mesocosm warming experiment. Zooplankton abundance and biomass was higher in mesocosms with larvae of heated origin than unheated origin, despite no difference in fish survival or chlorophyll *a* concentration. As fish larvae presence resulted in lower zooplankton abundance and biomass than in mesocosms without fish and the removal of most large-sized zooplankton (Supplement: Fish predation), the origin effect on zooplankton was likely due to feeding variation in fish. Zooplankton biomass data indicated that perch larvae originated from the heated area generally feed less than the ones of unheated origin, especially when experimental temperature exceeded the natural water temperature. Our results suggest that perch may have responded to the long-term extensive heating in ways that resulted in a reduced feeding on zooplankton during their larval stages and that multi-generational heating-induced changes in predators can have significant impact on their prey communities.

The reduced top-down effect of larvae of heated origin on the zooplankton communities, compared to that of larvae of unheated origin, might have arisen from a few non-exclusive mechanisms. Firstly, perch from the chronically warmed environment may have physiologically adapted such that they require less food to sustain themselves under elevated temperatures due to a lowered metabolic rate (14,58). If this is the case, it could enable high growth rates also at a lower feeding rate by reduced energy spent on maintenance resulting in a higher energetic efficiency (i.e. difference between energy intake and metabolic costs). In the populations that our larval fish originated from, adult perch from the heated habitat display significant metabolic thermal compensation and lowered heart beat at high temperatures compared to adults from the unheated population (35). Such warming-induced metabolic changes are commonly due to phenotypic plasticity (59), but there is also evidence showing that changes in metabolism of fish can reflect evolutionary adaptations, potentially to compensate for higher energy losses at high temperatures (14). Secondly, reduced attack rates can directly decrease top-down effect. When exposed to high temperatures, some organisms display reduced attack rates also after acclimatization to warmer environments (5,58). Thirdly, shifts in growth and development rates can be linked to evolutionary changes in response to warming (36). As a (potentially) higher energetic efficiency may also have helped fish maintain high growth rates, this might partly explain why larvae of heated origin gained slightly yet significantly more weight compared to larvae of unheated origin. Young (though non-larval) perch from the heated area also have a faster growth rate and an energy allocation strategy for earlier reproduction (20,21). Ultimately, these changes can lead to changes in body size, which also may result in differences in their diets (39,60). However, body size at inoculation and the end of experiment of larvae did not differ significantly between the thermal origins in our experiment. It is also possible that perch of heated origin have adapted to their already warm-adapted new prey communities and thus fed differently when exposed to a zooplankton community from the unheated area. While more comprehensive testing for the genetic base of local adaptation due to warming is needed, microsatellite DNA analysis of roe strands collected for our experiment already showed low but significant population differentiation between the thermal origins, which may partly be driven by adaptation to the higher temperatures. The results of our mesocosm experiment demonstrated that such impacts of chronic warming across multiple generations on wild fish populations can also affect their prey communities.

In addition to the higher zooplankton abundance, there were proportionally less large-sized zooplankton in mesocosms with larvae of heated origin than in those with larvae of unheated origin at high temperatures, and vice versa at low temperatures (Figure 3). This difference in zooplankton size- and species-composition could be due to different feeding preferences between perch larvae depending on thermal origin. Perch larvae of heated origin might have grown faster at higher temperatures, and thus having a larger gape size (than larvae of unheated origin) enabling them to predate more on larger-sized zooplankton (61). This may result from adaptation to long-term heating (20,27) and if heritable, provide an advantage for offspring in coping with a predictable chronically warm environment. This might partly explain the somewhat larger weight increase of larvae of heated than of unheated origin.

We found that the larvae origin effect on zooplankton abundance and composition was most pronounced at the highest end of the experimental temperature range (25 °C), which is a temperature that the chronically heated environment frequently reaches during the perch hatching season (Figure S1). This interaction between experimental temperature and larvae thermal origin suggests that the consequences of the potential adaptation in fish induced by the long-term heating can be temperature specific, as changes due to warming seem to modify fish top-down effects. This might have occurred due to the temperature-dependence of attack rates or handling time (5,11,40,62). In our experiment, the zooplankton prey responses to fish predation varied with the experimental temperature and was most pronounced when the experimental temperature reached the natal temperature of the larvae of heated origin.

However, the effect of fish thermal origin on zooplankton did not cascade down to affect phytoplankton biomass, as we found no differences in chlorophyll *a* concentration between mesocosms containing larvae of different origins (Figure S10). Primary producers might not be as susceptible to moderate changes in the feeding of top predators as the intermediate trophic levels (12) because the effect is mediated by trophic interactions across several trophic levels when it cascades through the food web. As we lack information about phytoplankton community composition, we are not able to rule out responses in phytoplankton composition depending on fish thermal origin. In the wild, primary producers would often be exposed to the same long-term environmental changes (e.g. warming) as higher trophic levels, such as fish, and would thus likely adapt in parallel (40), but we set up our experiment with prey communities of only natural (unheated) origin and not with separate treatments of the prey community of heated or unheated origin.

Besides changes in fish larvae feeding due to potential warming adaptation, the effect of fish thermal origin on the experimental zooplankton communities could also be explained by maternal effects from fish being acclimatized to different thermal regimes. Despite our effort to control for such differences, the roe strands collected from the heated area (5.8 ± 1.2 cm) were larger in width than the ones collected from the unheated area (3.9 ± 0.4 cm; Welch Two Sample t-test p < 0.05, effect size 0.6). Perch roe strand width is generally correlated with larger body size at hatching (61). This may explain why, despite hatching around the same time (May 18^th^), body lengths of larvae of heated origin were slightly larger than larvae of unheated origin at inoculation (Welch Two Sample t-test p < 0.05, effect size 0.5), however, no difference was found in larvae body weight. We argue that any maternal effects and acclimation before roe collection were likely small as egg strands were kept until hatching in a single hatchery for a much longer period with uniformed conditions and variation in all other environmental factors was minimized.

Our mesocosms were open to the atmosphere, making the experimental water temperatures vary with weather conditions, but with consistent temperature differences between mesocosms. On the other hand, our field conditions may be more relevant to climate change effects in nature than studies conducted at constant temperature as the latter have proved to be inaccurate at predicting responses to fluctuating conditions in ectotherms (63). We did not systematically quantify the presence of other organisms besides zooplankton in the mesocosms, however, we detected presence of chironomids larvae in 26 out of 38 mesocosms from spawning midges, zooplankton eggs at very low densities plus that we have observed zooplankton taxa emerge in the middle of the experiment. These organisms could have contributed to the larval fish diet without being accounted for in our assessment of the prey community. Changes in fish feeding may thus have not fully have been reflected in the sampled zooplankton community due to additional food sources that we did not sample. This could have contributed to variation in response variables among mesocosms within treatments. However, more than 99% of the fish larvae in the mesocosms were smaller than 20 mm and thus likely unable to efficiently consume larger prey such as chironomid larvae or pupae (64). Similarly, zooplankton eggs and protists not accounted for are likely also of minor importance given larval perch feeding preferences (65). Thus, we believe the observed differences in zooplankton prey communities are driven by fish larvae thermal origin.

There are also trophic interactions within the zooplankton community that may influence the results. Among species detected in our mesocosms, some meso-zooplankton species (e.g. *Cyclops* sp.) may prefer to predate on micro-zooplankton (20-200 μm) instead of merely grazing on phytoplankton (66). Fish predation on predatory zooplankton species can therefore result in both competitive and potentially predatory release on rotifers, which might explain the high numbers (>1000 individuals, taking up more than 70% of all individuals) of *Keratella* species in some mesocosms towards the end of the experiment. Besides by direct removal through predation, presence of fish larvae might create a landscape of fear for zooplankton that indirectly affect zooplankton community composition and dynamics (26,67). Measures on behavioural or chemical cues were, however, out of the scope of our experiment but may be informative in future experiments on indirect effects of potential adaptation to warming in aquatic predators.

In conclusion, we show that cross-generation changes in wild predators in response to whole-ecosystem warming can have significant cascading effects on prey communities via predator feeding behaviour. Although warming can change predator feeding across generations (11), ecosystem effects may depend on whether trophic interactions buffer or amplify the cascading effects of warming. Thus, it is important to measure direct responses in the prey community.

We therefore call for experiments generalizing our novel findings, to test whether responses to multi-generational warming in other ectothermic predators and/or older life stages would cascade down to affect other trophic levels and to further investigate the mechanisms by incorporating evolutionary and quantitative genetic methods – to infer whether these responses to warming are due to local adaptation, maternal effects or others.

## Acknowledgement

We are grateful to Carolina Åkerlund and Fredrik Landfors for the help during perch roe strand collection, Olivia Bell for assisting in the experiment, Wojciech Uszko for the generous guidance and directions on zooplankton identification, calibration and analysis, the Functional Microbial Ecology Lab (FUME) at SLU for facilitating the chlorophyll sample processing, and Oksana Burimski at Estonian University of Life Sciences, Institute of Veterinary Medicine and Animal Sciences, Chair of Aquaculture for microsatellite DNA processing.

## Ethics

This experiment was conducted in accordance with national regulations for animal care, and the experimental design and practices were reviewed and approved by the regional review board for ethical animal experiments in Uppsala, Sweden. Approved permit number: Dnr 5.8.18-04546-2021, permit holder AG.

