## Supplementary_materials for "Multi-decadal warming alters predator’s effect on prey community composition"

### Perch larvae thermal origin

**Figure S1**. Map of study area (a, zoomed in b) and daily water temperature (c) in the heated vs. unheated area throughout the year, showing a > 5^◦^C temperature difference between the areas. The red and blue arrows in subplot b indicate heated and natural temperature water flow direction underwater through pipes. The “Heated” marked the enclosed ecosystem. The white crosses show where the perch roe strands were sampled. The daily mean water temperature measured is shown as red (heated) and blue (unheated) opaque points. The thick red and blue lines mark the maximum temperature in the heated and unheated area. Water temperature was measured with temperature loggers during the ice-free period (March-December; 1989-2003; only including temperature measured at the same depth ~ 0.5 m and time for both areas). The more solid the color, the more data points with that temperature measured were included.

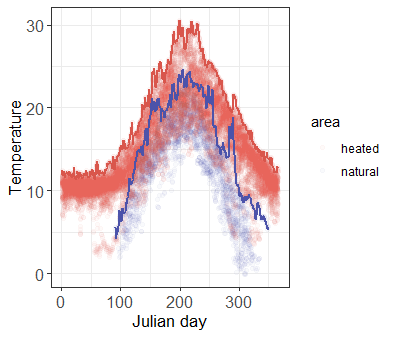

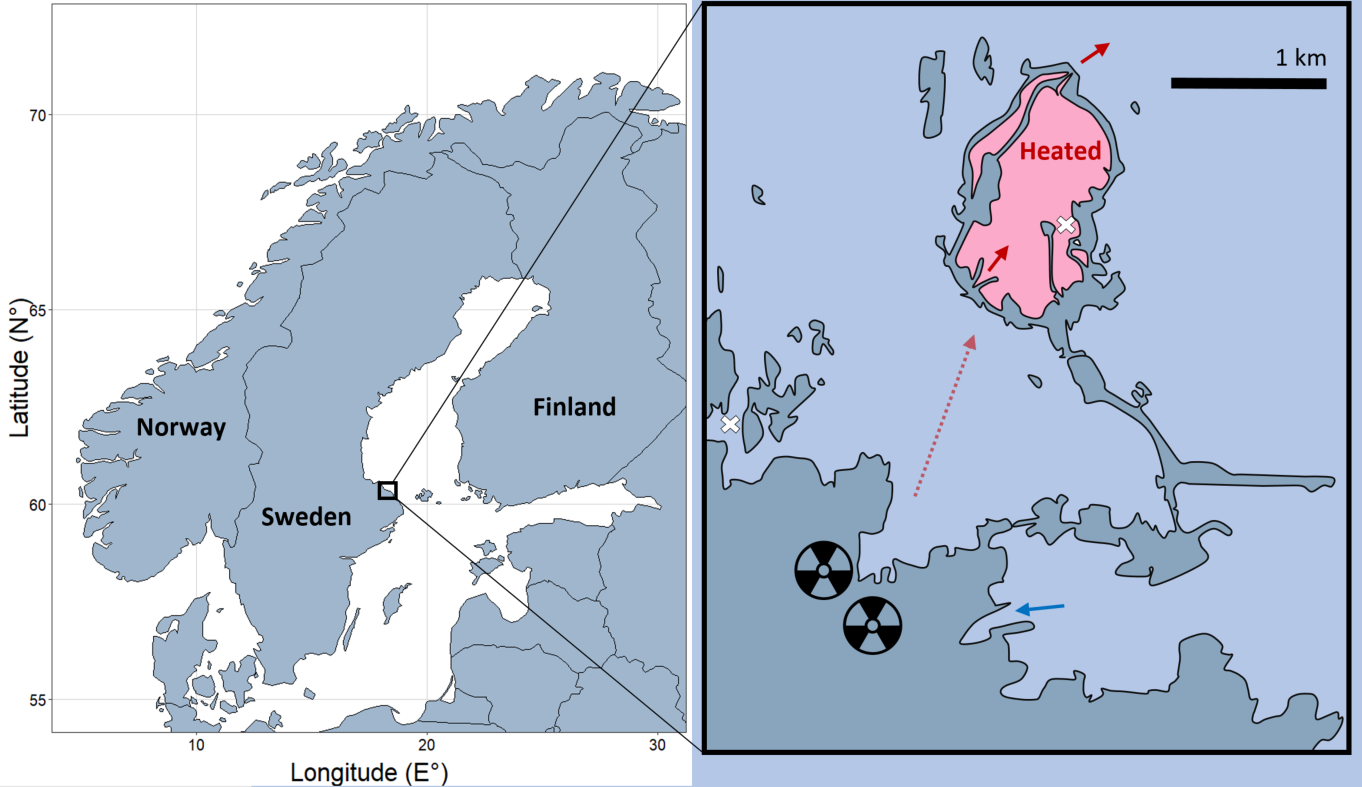

**a**

**b**
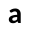

**c**
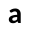

#### Similarities and differences between areas

Except for the large difference (>5^◦^C) in water temperature described in Figure S1 (daily means of hourly measurements using temperature loggers, across the years 1989-2003), the two areas -from which the perch larvae used in the experiment originated from- share many similarities: air temperature, weather conditions and any long-term environmental trends subjected to the whole region, such as climate change. Because the heated area was constructed from connecting existing small islands in the archipelago, and the unheated area is part of the same archipelago, they also share the same general coastal habitat features such as being shallow and near-shore. As water in the heated area has been collected from the natural temperature area (indicated by the blue arrow, Figure S1), pumped through the power plant to absorb heat and then exiting into the heated area via pipes (red arrow with dashed line, Figure S1), the two areas most likely share similar water quality as it is the same water circulating within this 3 km radius area of the same archipelago, except for the flow-through current. Available water quality data that was monitored the same way throughout the study period is water transparency (measured as secchi depth). Sandström & Karås (2002) showed that the water transparency is approximately the same in both areas (“Ön” is our control area with secchi depth 3-5 meters and “Biotest basin” is our heated area with secchi depth 4-5 meters, Table 1 in Sandström & Karås 2002). And previous studies have reported no differences in salinity (Snoeijs & Murasi, 2004), light (Huss et al., 2021) or dissolved nutrients (Hillebrand et al., 2010) between the two areas, although it should be noted that for most years no data is available.

### Larvae hatching

Fifteen roe strands were collected from each of the heated and unheated area (Figure S1). From each area, eight roe strands with similar adhesion and width were selected and placed in separate 100 L aquariums, labelled as “Heated” (H) 1-8 and “Unheated” (UN) 1-8 (see attached Table S1.xlsx for details) and the other seven roe strands from each area were placed in 40 L aquaria. They were maintained at room temperature and on a 16h**:**8h light to dark schedule, and thus all subjected to the same temperature and light regimes.

### Experimental set-up

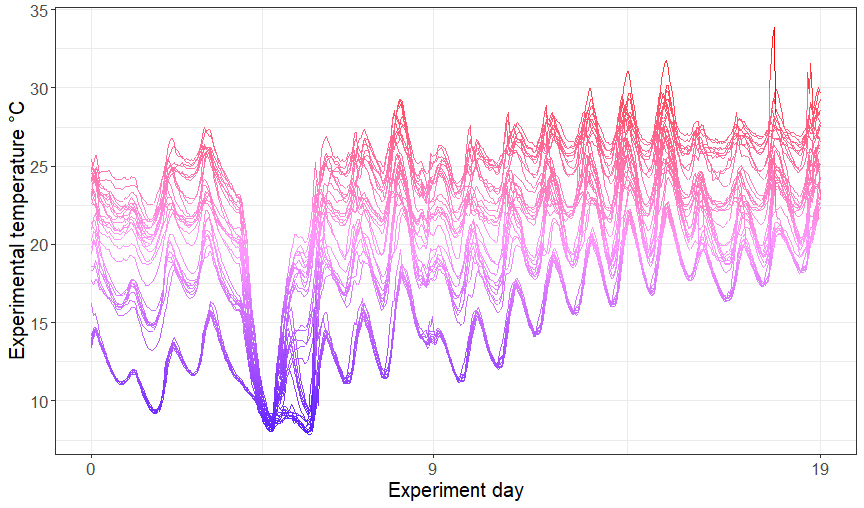

**Figure S3**. Hourly measured experimental temperature in each mesocosm shows temperature fluctuations between and within each mesocosm. Our mesocosms were open systems, which is why the experimental temperatures varied with weather conditions and diurnal cycles, but the variance between mesocosms were mostly consistent. However, a heavy rainstorm swept our experiment site on day 4-5, which led to a power failure that shut down some thermostats and air pumps. This resulted in a sudden drop in temperature and between-mesocosm variance.

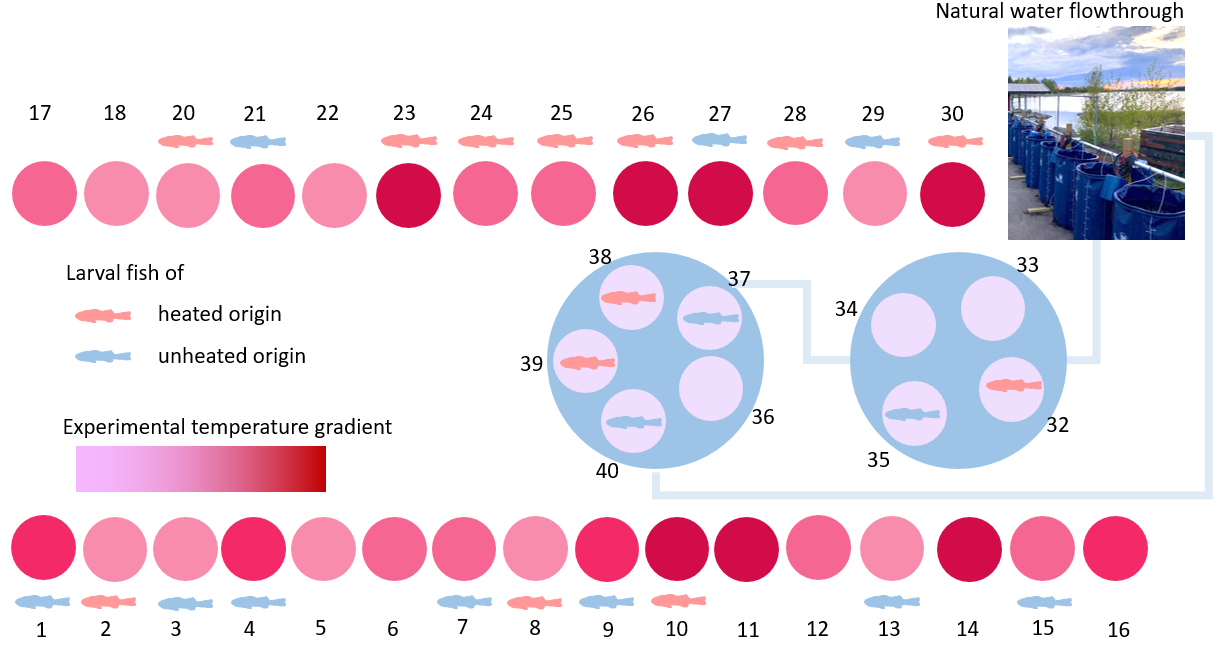

**Figure S2**. Mesocosm experimental set-up, with 38 outdoor tanks of which 26 were inoculated with fish larvae (13 of heated origin indicated by red fish, and 13 of unheated origin indicated by blue fish), and with the remaining 12 mesocosms kept without fish as a control. A thermal gradient of 14 - 25 °C was maintained among the 38 mesocosms. In 29 mesocosms thermostats were set to maintain 18, 22 and 26 °C and in 9 mesocosms, thermostats were inside the tanks but turned off. A water flow-through system with natural sea water surrounded the 9 mesocosm tanks placed in pools (mesocosm numbered 32-38) to keep the water temperature cooler than air.

**Table S2**. The larval and temperature treatment for each mesocosm, and numbers of **larval fish caught on experimental day 20 and** July 2^nd^. Experimental temperature (in °C) is the average of hourly measurements midwater from the beginning of the experiment until the respective sampling day. Heating started the day before Day 0.

| Mesocosm | Fish origin | Experimental temperature °C | | | Number of fish caught | | |
| --- | --- | --- | --- | --- | --- | --- | --- |
|  |  | Day 0 | Day 9 | Day 19 | Day 20 | July 2^nd^ | Total |
| 1 | Unheated | 22.35 | 21.1 | 23.82 | 2 | 0 | 2 |
| 2 | Heated | 17.73 | 16.58 | 19.08 | 8 | 0 | 8 |
| 3 | Unheated | 17.19 | 16.7 | 19.1 | 9 | 0 | 9 |
| 4 | Unheated | 19.23 | 20.78 | 23.2 | 9 | 1 | 10 |
| 5 | NA | 18.33 | 17.05 | 18.92 | / | / | / |
| 6 | NA | 22.15 | 20.4 | 21.87 | / | / | / |
| 7 | Unheated | 21.85 | 19.9 | 21.53 | 6 | 0 | 6 |
| 8 | Heated | 16.66 | 16.09 | 18.29 | 9 | 0 | 9 |
| 9 | Unheated | 22.81 | 21.71 | 23.97 | 1 | 0 | 1 |
| 10 | Heated | 22.95 | 21.85 | 24.53 | 2 | 1 | 3 |
| 11 | NA | 22.35 | 22.07 | 24.31 | / | / | / |
| 12 | NA | 20.5 | 19.23 | 21.21 | / | / | / |
| 13 | Unheated | 17.64 | 16.26 | 18.17 | 6 | 3 | 9 |
| 14 | NA | 22.5 | 21.9 | 24 | / | / | / |
| 15 | Unheated | 21.55 | 20.32 | 21.89 | 3 | 1 | 4 |
| 16 | NA | 22.23 | 21.52 | 23.45 | / | / | / |
| 17 | NA | 21.55 | 20.02 | 21.82 | / | / | / |
| 18 | NA | 19.66 | 17 | 19.04 | / | / | / |
| 20 | Heated | 17.54 | 17.01 | 19.29 | 9 | 1 | 10 |
| 21 | Unheated | 21.92 | 21.1 | 22.29 | 5 | 2 | 7 |
| 22 | NA | 18.23 | 17.52 | 19.38 | / | / | / |
| 23 | Heated | 22.55 | 22.1 | 24.97 | 0 | 0 | 0 |
| 24 | Heated | 21.18 | 20.19 | 21.4 | 5 | 1 | 6 |
| 25 | Heated | 21.07 | 20.53 | 21.63 | 6 | 0 | 6 |
| 26 | Heated | 21.87 | 22.61 | 24.48 | 2 | 1 | 3 |
| 27 | Unheated | 21.9 | 22.34 | 24.22 | 0 | 2 | 2 |
| 28 | Heated | 21.76 | 20.11 | 21.7 | 8 | 0 | 8 |
| 29 | Unheated | 16.83 | 16.36 | 18.22 | 9 | 1 | 10 |
| 30 | Heated | 23.62 | 22.85 | 24.87 | 2 | 1 | 3 |
| 32 | Heated | 13.32 | 12.55 | 15.15 | 7 | 1 | 8 |
| 33 | NA | 12.91 | 12.09 | 14.72 | / | / | / |
| 34 | NA | 12.39 | 12.16 | 14.78 | / | / | / |
| 35 | Unheated | 12.47 | 12.23 | 14.88 | 10 | 0 | 10 |
| 36 | NA | 12.96 | 12.16 | 14.82 | / | / | / |
| 37 | Unheated | 12.95 | 12.21 | 15 | 8 | 0 | 8 |
| 38 | Heated | 12.53 | 12.3 | 15.01 | 1 | 0 | 1 |
| 39 | Heated | 12.39 | 12.28 | 14.98 | 7 | 0 | 7 |
| 40 | Unheated | 12.58 | 12.31 | 14.99 | 4 | 0 | 4 |

### Mesocosm initial conditions

#### Phytoplankton

Phytoplankton community biomass was approximated by estimation of chl *a* concentration (µg/L) in our mesocosms (Huot et al., 2007). Chlorophyll *a* concentration in each mesocosm was derived from a chl *a* calibration curve (with a linear relationship, Equation 1). Equation 1 was obtained from measured fluorescence of solutions with a gradient of chl *a* concentrations. We measured fluorescence (RFU) using a Hidex Sense microplate reader of 18 different solutions of a range of chl *a* concentrations (0-500 µg/L) diluted from standard stock solution (10mg/L).

Chl *a* concentration (μg/L) = (RFU - 2766.5)/194.1 (Equation S1)

There were no significant differences in chlorophyll *a* concentration (sampled on day -1) among mesocosms with fish of different origins or without fish: ANOVA, chl *a* F(2,35) = 0.859, P = 0.432.

#### Zooplankton

There were no significant differences in zooplankton abundance (sampled or zooplankton biomass (day 0) among mesocosms with fish of different origins or without fish: ANOVA, zooplankton abundance F(2,32) = 0.451, P = 0.641, zooplankton biomass F(2,32) = 0.231, P = 0.795). However, temperature seems to already have an effect at day 0 on the zooplankton abundance, F(1,32) = 16.256, P = 0.0003 although a small negative effect (estimate from linear relationship = -0.09685) with higher zooplankton abundance in cooler mesocosms. Nevertheless, zooplankton communities were introduced only one day before the experiment started, how much temperature could affect the abundance during that one day is arguable. The same goes for chl *a*, which was sampled two days prior to the experiment start, without temperature treatment in place.

**Table S3**. Assumed length-to-weight relationships for zooplankton, where $W$ is dry weight in µg, and $L$ is body length in mm (prosome length for copepods, and the longest body axis excluding spines for all other taxa). All relationships are based on Bottrell et al. 1976.

| Taxon | Conversion formula |
| --- | --- |
| Copepoda (incl. Nauplius) | ln W = 1.9526 + 2.399 * ln L |
| Keratella quadrata, K. cruciformis | ln W = ln(92.224) + 2.955 * ln L |
| Keratella cochlearis | ln W = ln(28.985) + 2.955 * ln L |
| Bdelloida | ln W = ln(16.949) + 3.0089 * ln L |
| Podon, Chydorus | ln W = 1.7512 + 2.653 * ln L |
| Bosmina | ln W = 3.0896 + 3.0395 * ln L |

Total biomass per zooplankton taxa in each sample was calculated as biomass_total_ = (m_1_ + m_2_ + … m_n_)/n_1_ * n_2_, where m_n_ is the mass (µg) of each of the n length-measured zooplankton individuals of that taxa, n_1_ is the total number of zooplankton of that taxa with converted mass and n_2_ is the number of individuals of the taxa counted in the sample.

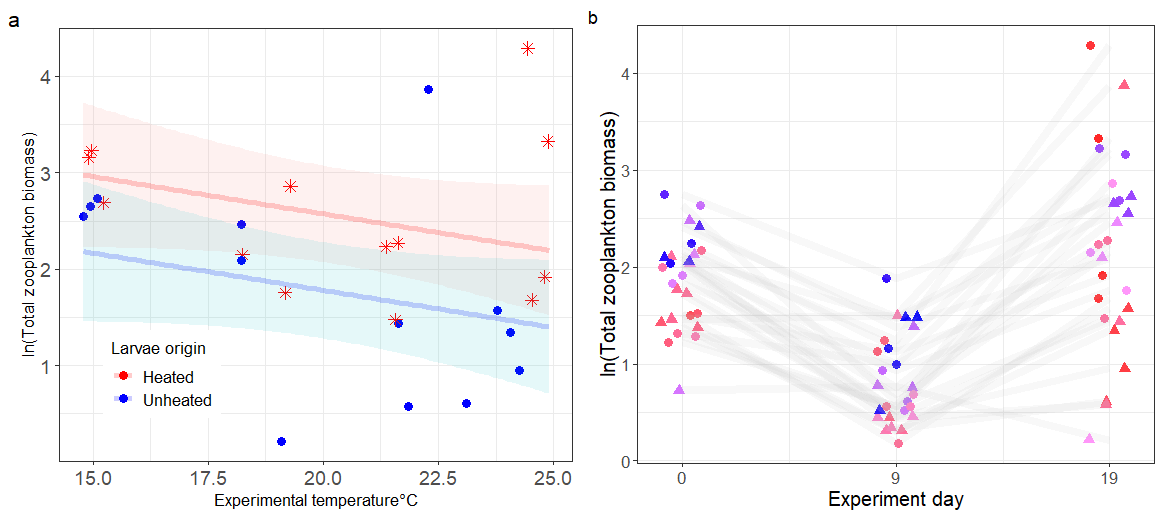

**Figure S4.** Zooplankton total biomass (µg) at the end of the experiment (a) and throughout the experiment (b) in mesocosms containing larval fish that originated from the heated (stars) or unheated area (circles) at different experimental temperatures. The red and blue lines depict the predicted ln-transformed zooplankton biomass and the belts show their corresponding confidence intervals at 95%. The effect of larvae thermal origin on zooplankton abundance is predicted to increase with temperature. Note, all mesocosms were measured on the same days (0, 9, 19), but have been displaced for visibility.

### Larval fish analysis

#### **Survival, length and weight**

**Figure S5**. (a) Total number of fish larvae caught at the end of the experiment (day 20, after the last zooplankton sampling on day 19), their (b) weight and (c) length. The number caught indicates that survival of larvae do not differ significantly (p > 0.05, ANOVA test) depending on larval thermal origin, but varies with experimental temperature (ANOVA, F(1, 22) = 6.61, P = 0.017 η2 = 0.23).

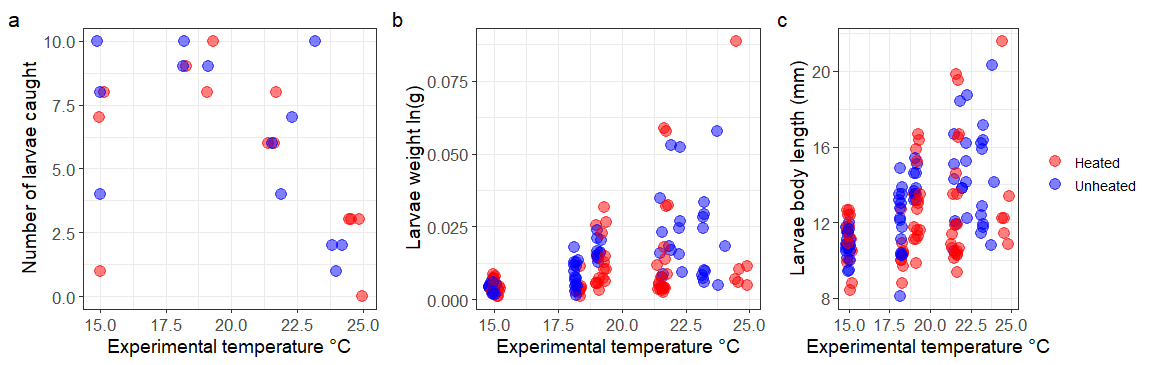

Larval fish survival of each mesocosm was equalled to number of fish caught (Figure S5a). Based on square root transformed catch data analysed with ANOVA (F(1,22) = 0.44, P = 0.48), there was no significant difference in survival among origins. However, larvae survival decreased with temperature, ANOVA, F(1, 22) = 6.61, P = 0.017, η2 = 0.23.

**In the attached Table S4.xlsx, we show wet weight, total length and** length increment **(**L_day20_ – L_inoculation_**) of larval fish of different origin caught in the mesocosms on the final experimental day (day 20), and upon emptying the mesocosms on** July 2^nd^ **(22 days after the experiment ended). Length increments are not included in any analysis for larvae caught on** July 2^nd^ **as the values are not comparable with larvae caught on day 20. We did not calculate weight increment because wet weight at inoculation was so low (two orders of magnitude lower) that it is negligible comparing to W**_day20_. On average, at the start of the experiment, a larva of heated origin weighed 0.428 ± 0.005 mg and a larva of unheated origin weighed 0.413 ± 0.003 mg compared to 11.388 ± 15.474 mg and 13.056 ± 12.183 mg, respectively, at the end of the experiment. Larvae from the heated origin grew to be heavier than those of unheated origin (Figure S5b). Temperature has a significant impact on larvae survival – survival decreased with temperature, ANOVA, F(1, 22) = 6.61, P = 0.017, η2 = 0.23, and individual weight F(1, 134) = 55.54, P < 0.0001, η2 = 0.28) as well as larvae body length (Figure S5c, length: F(1, 134) = 41. 17, P < 0.0001, η2 = 0.23) increased with temperature.

**Table S5**. Weight and total length of perch larvae in aquaria measured at the start of the experiment (Day 0) into the mesocosms. Wet weight was measured in groups of five to increase precision. “/” marks individuals that could not be measured due to freeze damage. Egg strand number corresponds to the ones in Table S1.

| larvae egg strand | weight of 5 individuals (g) | average weight (g) | average total length (mm) |
| --- | --- | --- | --- |
| heated1 | 0.0024 | 0.00048 | 5.86 |
| heated2 | 0.0021 | 0.00042 | 5.56 |
| heated4 | 0.0018 | 0.00036 | 5.38 |
| heated5 | 0.0021 | 0.00042 | / |
| heated8 | 0.0023 | 0.00046 | 5.75 |
| unheated1 | / | / | 5.85 |
| unheated3 | / | / | 5.66 |
| unheated4 | 0.0022 | 0.00044 | 5.71 |
| unheated6 | 0.0021 | 0.00042 | 4.38 |
| unheated7 | 0.0019 | 0.00038 | / |

As for roe strand widths, those of heated origin were slightly larger (5.8 ± 1.2 cm) than those from the unheated area (3.9 ± 0.4 cm). At hatching, larvae of the heated origin were also larger on average (5.72 ± 0.42 mm) than the larvae of unheated origin (5.61 ± 0.39 mm; Welch Two Sample t-test p < 0.001, effect size glass’s delta 0.5). Larval weight at hatching, however, showed no difference between larvae thermal origins (Table S5, p = 0.609, glass’s delta 0.5). Note, however, that individuals were measured in groups of five as they were too light and too soaked in ethanol to weigh individually.

These roe strands were likely maximum 10-day post spawning, based on water temperature measurements and the absence of visible embryos at the time of collection (Wang & Eckmann, 1994). There was no difference in temperature between hatching aquaria (Welch Two Sample t-test, P = 0.94, effect size in Cohen’s d = 0.014) and the roe strands started hatching around the same time irrespective of origin. The first roe strand from the heated area started hatching on May 17^th^, all roe strands had started hatching on May 18^th^ and more than half of each roe strand had hatched on May 21^st^. The first eggs from the unheated area hatched on May 18^th^, all had started to hatch on May 19^th^ and more than half of each strand had hatched on May 21^st^ (hatching record, Table S1).

#### Genetic analysis

The microsatellite data was analyzed using Fisher’s exact probability test via Genepop version 4.7.5 (<https://genepop.curtin.edu.au/>) options Population Differentiation and Fst & other correlations. The genotypes based on the 14 microsatellite loci selected can be found in the attached 14_microsatellite_primer.xlsx and microsatelliteDNA_genotypes.xlsx.

### Zooplankton biomass and composition

**Table S6.** Results of model selection for the zooplankton biomass based on model pair testing - likelihood ratio tests (significance level between the new model and null model). Model pairs are: null and second, second and third, and so on. We judged whether including larval origin, or experimental temperature, or larval origin × temperature, improved the model explaining the total variance in zooplankton biomass based on both the significance of likelihood ratio tests and goodness of fit measures. The coefficients of the best models (in bold) are: origin (unheated) -0.79, temperature -0.077 and origin (unheated): -0.47, temperature: -0.059. * indicates significance level.

|  | End of experiment | | Throughout the experiment | |
| --- | --- | --- | --- | --- |
|  | Formula | Significance | Formula | Significance |
| ln(total zooplankton biomass + 1) ~ | 1 |  | day + random(day\|mesocosm) |  |
|  | temperature | 0.067* | temperature + day + random(day\|mesocosm) |  |
|  | **origin + temperature** | **0.028*** | **origin + temperature + day +random(day\|mesocosm)** | 0.076 |
|  | origin × temperature | 0.25 | origin *×* temperature + day +random(day\|mesocosm) | 0.519 |

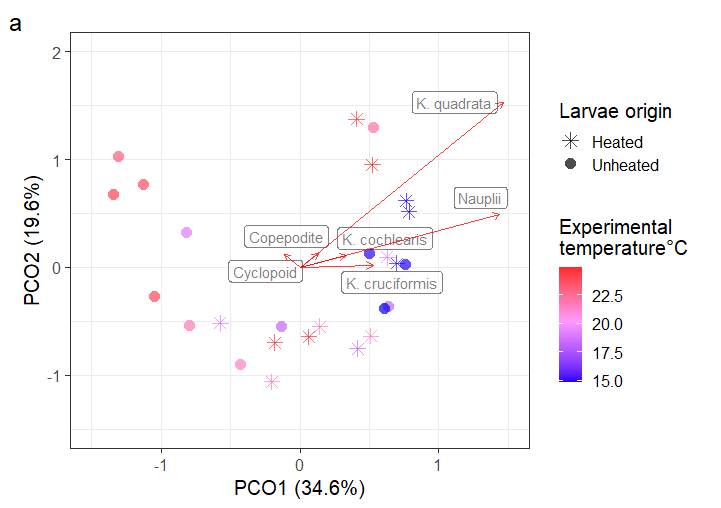
**Figure S6**. (a) Ordination showing PCO1 (explaining 34.6% of the total variation) and PCO2 (explaining 19.6%) results on zooplankton community composition based on biomass of different taxa, on experiment day 19 in mesocosms with larvae of heated (star) or unheated (circles) origin. Colour shows mean experimental temperature from cold (blue) to warm (red). The red arrows show the six taxa having the highest absolute scores on PCO1. (b) Predicted PCO1 at given experimental temperatures, shown by solid lines and 95% confidence intervals shown by the
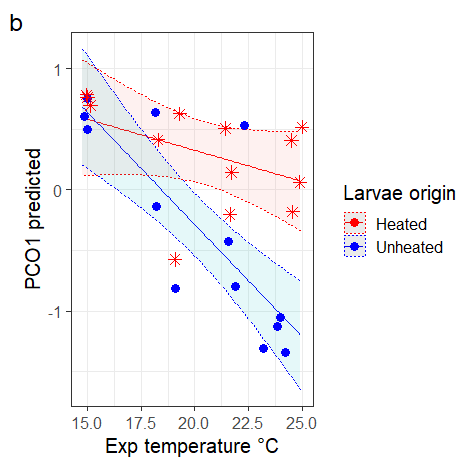
dashed lines. The difference in zooplankton composition between larvae origins increased with experimental temperature indicated by ANOVA results: on temperature F(1,22) = 19.685, P = 0.00021; on origin F(1, 22) = 13.40, P = 0.00138 and on interaction between temperature and origin, F(1,22) = 7.20, P = 0.0136. The six most influential taxa are the same as for zooplankton abundance.

**Figure S7.** Zooplankton species composition shown by relative abundance on day 0, 9 and 19 in each mesocosm with larvae of heated origin (red fish) or unheated origin (blue fish). The mesocosms (numbered columns) are ordered from left to right following the lowest experimental temperature to the highest. Mesocosms with “unheated” larvae have proportionally more nauplii and copepods at high temperatures on day 19.

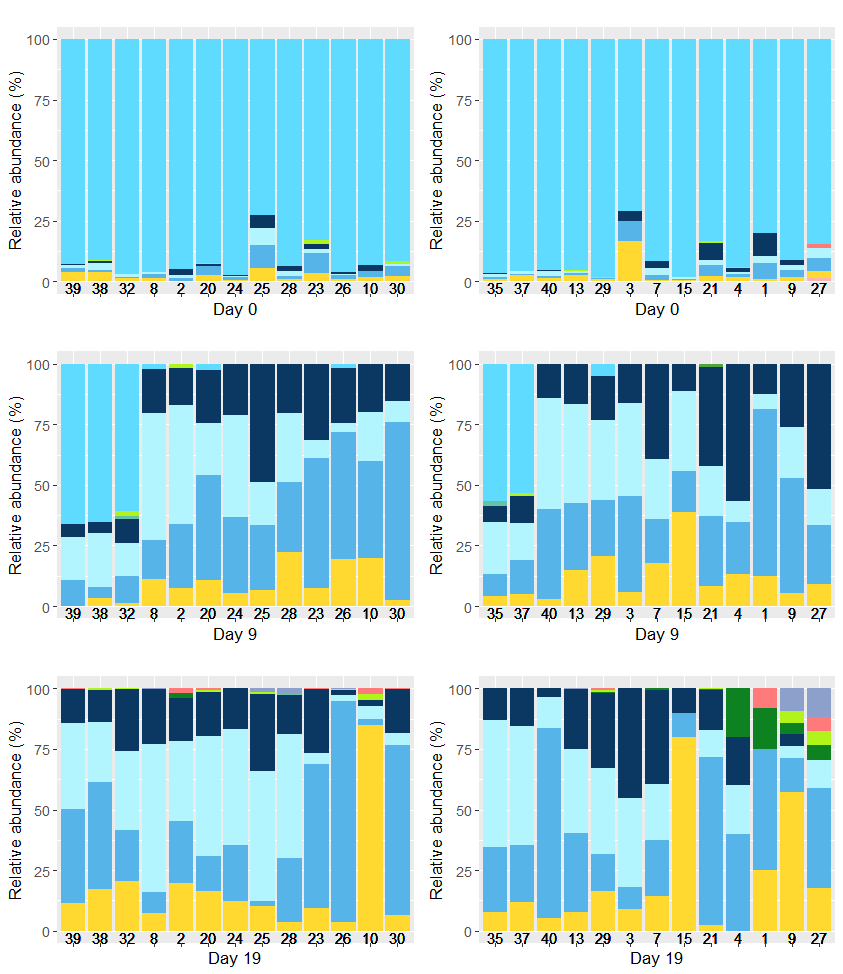

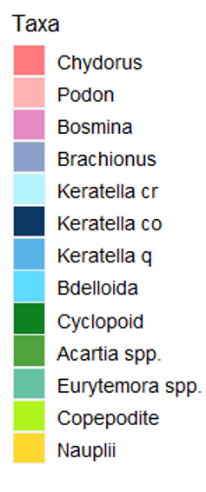

**Table S7**. Results of model selection on which model best explain the proportion of large-sized zooplankton, P(large zooplankton), based on model pair testing - likelihood ratio tests (significance level between the new model and null model). Model pairs are: null and second, second and third, and so on. Model in bold is the selected best model and * indicates significance level. On the right are the coefficients of the best model.

|  | Model selection | | **Best model:** **P(large zooplankton) ~ origin × temperature** | | |
| --- | --- | --- | --- | --- | --- |
|  | Formula | Significance | Term | Significance | Estimate |
| P(large zooplankton) ~ | temperature | 0.07 | Origin | < 0.001***** | -9.44 |
|  | origin + temperature | 0.83 | Temperature | 0.24 | -0.07 |
|  | **origin × temperature** | **< 0.001*** | Origin × temperature | < 0.001***** | 0.46 |

### Mesocosm water chemistry

We measured dissolved oxygen (DO), salinity and pH using a portable multi-parameter Aquaprobe (AP 2000, AquaReadLtd, UK). DO measurement was calibrated with a 2-points method: 0% with zero oxygen tablets (Mettler Toledo) and 100% using wet paper towel surrounding the probe (following AquaReadLtd’s manual instruction). The pH measurement was calibrated with a 3-points method using pH buffers at 7, 4 and 10. DO concentration decreased with temperature (F(1, 32) = 355.92, P < 0.0001) but the range was narrow: 8.32 – 9.40 mg/L and oxygen was saturated across the temperature gradient (≥ 100%). DO did not differ between mesocosms with fish of heated or unheated origin or no fish (F(2,32) = 1.524, P = 0.233). Salinity (PSU) at the end of the experiment ranged from 4.55 to 5.50 PSU, with no difference among mesocosms depending on whether there were fish present or fish origin (ANOVA, F(2,32) =0.602, P = 0.554). Salinity, however, increased with temperature, F(1,32) = 127.12, P < 0.0001. Water pH did not show any statistical significant difference depending on larval thermal origins, or fish presence/absence (F(2,32) = 3.031, P = 0.0623) or temperature (F(1,32) = 3.548, P = 0.0687), but there were a trend of pH increasing with temperature and being higher in mesocosms with larvae of heated origin.

a

b
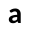

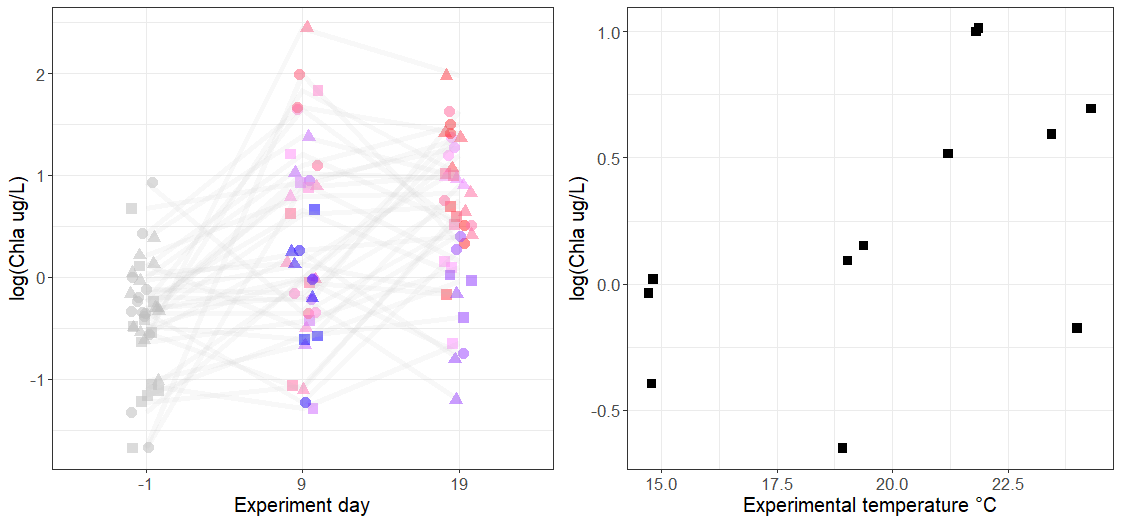

**Figure S8**. The left graph shows the natural log transformed chl *a* concentration in μg/L in mesocosms with larvae of heated (circles) or unheated (triangle) origin and mesocosms without fish (squares) throughout the experiment. There colours indicate the experimental temperature, from cold (blue) to warm (red). As temperature treatments were not in place on day -1, chl *a* concentration on that day is shown in grey. The right graph shows the natural log transformed chl *a* concentrations in mesocosms without fish on experiment day 19.

No difference was found in chl *a* concentration between mesocosms with larvae of heated or unheated origin (ANOVA, F(1, 72) = 1.10, P = 0.298). Concentration of chl *a* was higher in mesocosms with than without fish (ANOVA on throughout: F(1,108)=3.53, P = 0.0628; end: F(1, 34) = 7.81, P = 0.0085). Concentrations of chl *a* increased with temperature throughout the experiment regardless of fish presence (ANOVA F(1, 108) = 44.03, P < 0.0001), or thermal origin (F(1, 32) = 34.33, P < 0.0001).

### Fish predation

In support of the conclusion that differences in zooplankton communities were due to larvae thermal origin (and not non-larvae aspects), we found that zooplankton abundance was higher in mesocosms without fish than mesocosms with fish (ANOVA, P< 0.05) independent of experimental temperatures (ANOVA, P = 0.93). The relative abundance of copepods was higher without fish (Figure S9). In mesocosms without fish, zooplankton relative abundance per taxon changed throughout the experiment and was also affected by experimental temperature (Figure S9).

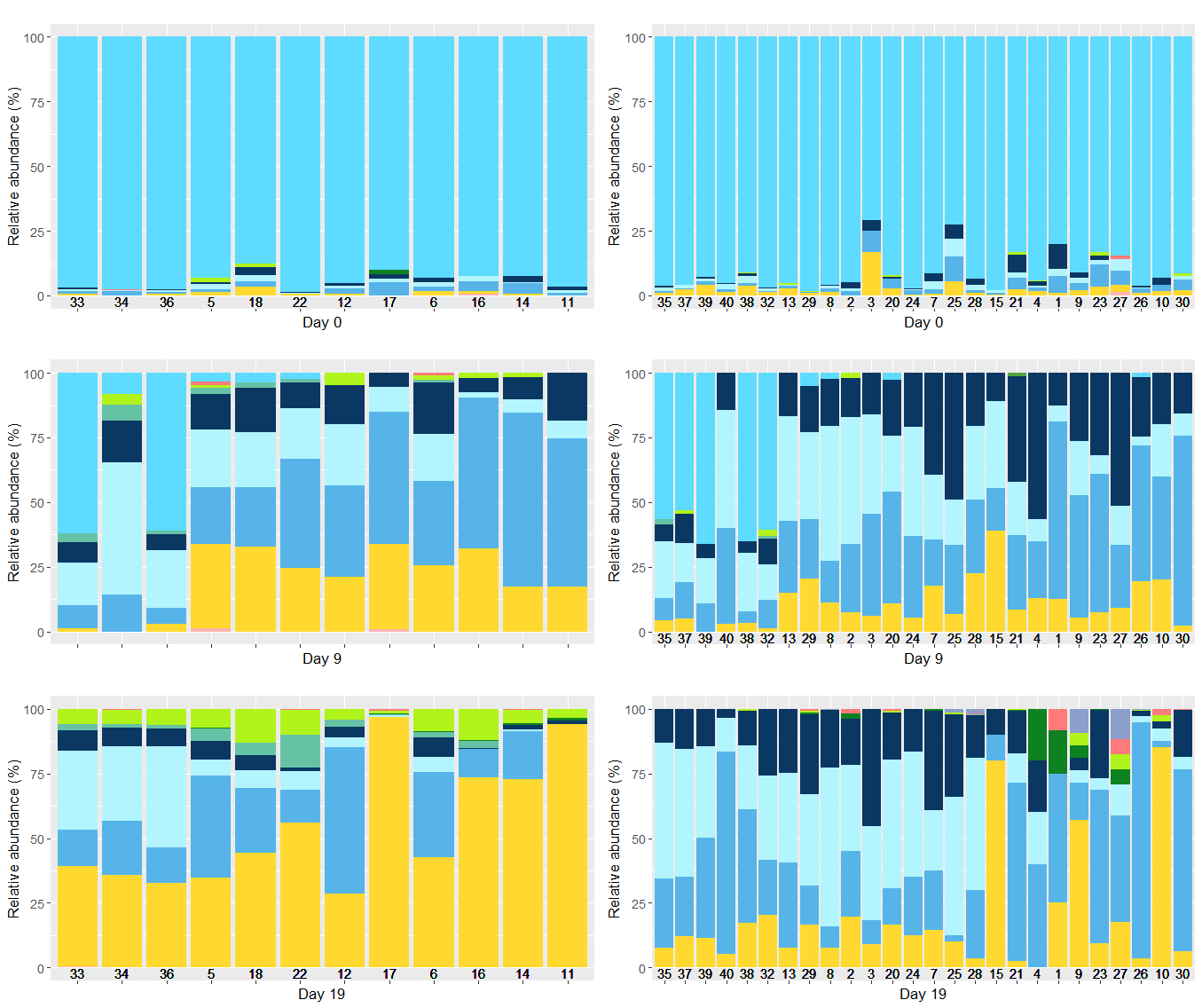

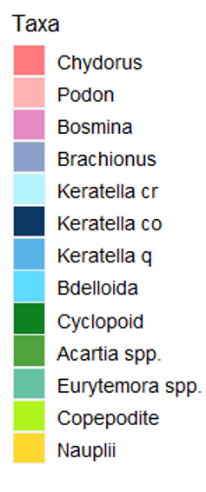

**Figure S9.** Zooplankton relative abundance per taxon shown as percentage on experimental day 0, 9 and 19 in mesocosms without fish (left) and with fish (fish silhouette). The numbered columns represent each mesocosm ordered from low to high temperature (left to right). Zooplankton community composition changed over time depending on whether there were fish or not. It also changed with temperature in both mesocosms with and without fish. The initially dominant taxon, order *Bdelloida*, was replaced by *Keratella* spp. and copepod nauplii. At the end of the experiment, copepod nauplii were dominant in some of the warmer mesocosms and more individuals of older stage copepodites, adult copepods and cladoceran emerged. Without fish, there were significantly more nauplii, copepodites and copopods at the end of the experiment.

### **Changes of plans**

We would like to share some of the hiccups we encountered during the experiment. The initial experiment design was to establish 40 mesocosms, with 4 replicates * 4 experimental temperatures * 2 larvae thermal origins (= 32 mesocosms) and 2 replicates * 4 temperatures in mesocosms without fish (= 8 mesocosms). Despite our extensive effort in mending the tanks from leaking with silicone, two leaked badly when filled with water. We therefore had to change the experiment design accordingly and thus ended up with 3 replicates * 4 temperatures * 3 fish treatments (2 thermal origins + 1 no fish) = 36 mesocosms, and assigned two additional mesocosms to the treatment with the highest temperature with fish.

While we set the thermostats at three fixed temperatures 18, 22 and 26 °C intending to keep three temperature levels (groups with similar variation), we ended up with a gradient of 14 - 25 °C. This is probably due to variation in heating between thermostats and placement of the mesocosms – some might have gotten more sunlight and some been more exposed to wind. So it would not be sensible to run statistical analyses with experimental temperatures as a factor with four levels.

**Out of 26 mesocosms with fish, we found 16 alive fish in 12 mesocosms when cleaning the tanks 22 days after the last day of the experiment. This shows that the method we implemented was not ideal and perhaps emptying all mesocosms would have been the best option.**
